# Global variations of Light Use Efficiency in Forests Jointly Driven by Plant Traits and Climatic Conditions

**DOI:** 10.64898/2026.06.16.732732

**Authors:** Yuan Zhang, Jie Lai, Yage Liu, Anzhi Wang, Xinghan Zhao, Jiabing Wu

**Author notes:** The journal (Ecosphere) provided us with many constructive suggestions, which inspired us to think more deeply. Considering the lengthy revision process, I’m uploading this manuscript to a preprint platform firstly.

## Abstract

Forests are essential to the global carbon cycle with light use efficiency (LUE) as a key parameter for assessing carbon sequestration capacity. However, the variations and drivers of LUE remain inadequately understood. Using remote sensing data, we analyzed global LUE patterns across five forest types and identified the main drivers. The global average annual LUE of forests is 0.93 ± 0.36 g C MJ^-1^ during the period 2001-2022, with an increasing trend of 0.0034 g C MJ^-1^ yr^-1^. Among forest types, evergreen broadleaf forests exhibited the highest LUE, followed by evergreen needleleaf forests. Deciduous broadleaf forests and mixed forests showed similar levels, while deciduous needleleaf forests exhibiting the lowest LUE. Variations in LUE were jointly driven by plant traits and climatic conditions, with generalized linear models explaining 86% and 98% of spatial and temporal LUE variations, respectively. These findings highlight the critical role of plant traits and climate in shaping forest LUE, providing insights for enhancing carbon cycle models and informing forest management strategies in the context of global change.

## 1. Introduction

Forests are a crucial component of the earth’s ecosystem, absorbing approximately 7.6 Gt CO_2_e yr^-1^ (Harris et al., 2021). In forest ecosystem, photosynthesis serve as the primary process for carbon fixation with light use efficiency (LUE) playing a direct role in determining its effectiveness. However, the pressure of global climate change, including the increasing frequency of extreme weather events (Kong et al., 2020; Robinson et al., 2021; Stott, 2016), alongside intensified human activities (Bourgoin et al., 2024; Zhou et al., 2023), have placed unprecedented strain on forest ecosystems. Therefore, understanding the spatial and temporal variations of forests LUE is essential for the accurately estimating gross primary productivity and the improving carbon cycle models.

LUE reflects how efficiently plants convert absorbed light energy into biomass (Monteith, 1972), serving as an important indicator of plant photosynthesis efficiency. It is widely used for estimating gross primary productivity through remote sensing, and various models built with this parameter are widely applied (Potter et al., 1993; Running et al., 2004; Wang et al., 2015; Xiao et al., 2004; Yan et al., 2015; Yuan et al., 2007). These models consider the light energy absorbed by the plants and the LUE under environment stress (Formula 1-3). Typically, each model assumes a theoretical maximum LUE value—representing optimal conditions—for various vegetation types, while climatic variables (e.g., temperature, soil moisture) limit this potential. Differences in LUE estimation methods are a primary distinction among these models (Table 1). However, a common issue arises with LUE utilization: many models apply fixed maximum values across vegetation types globally, overlooking spatial heterogeneity within similar vegetation types. This raises a critical question: can the LUE values derived from such assumptions truly represent the variability within vegetation parameters?

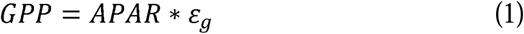

**Table 1.**
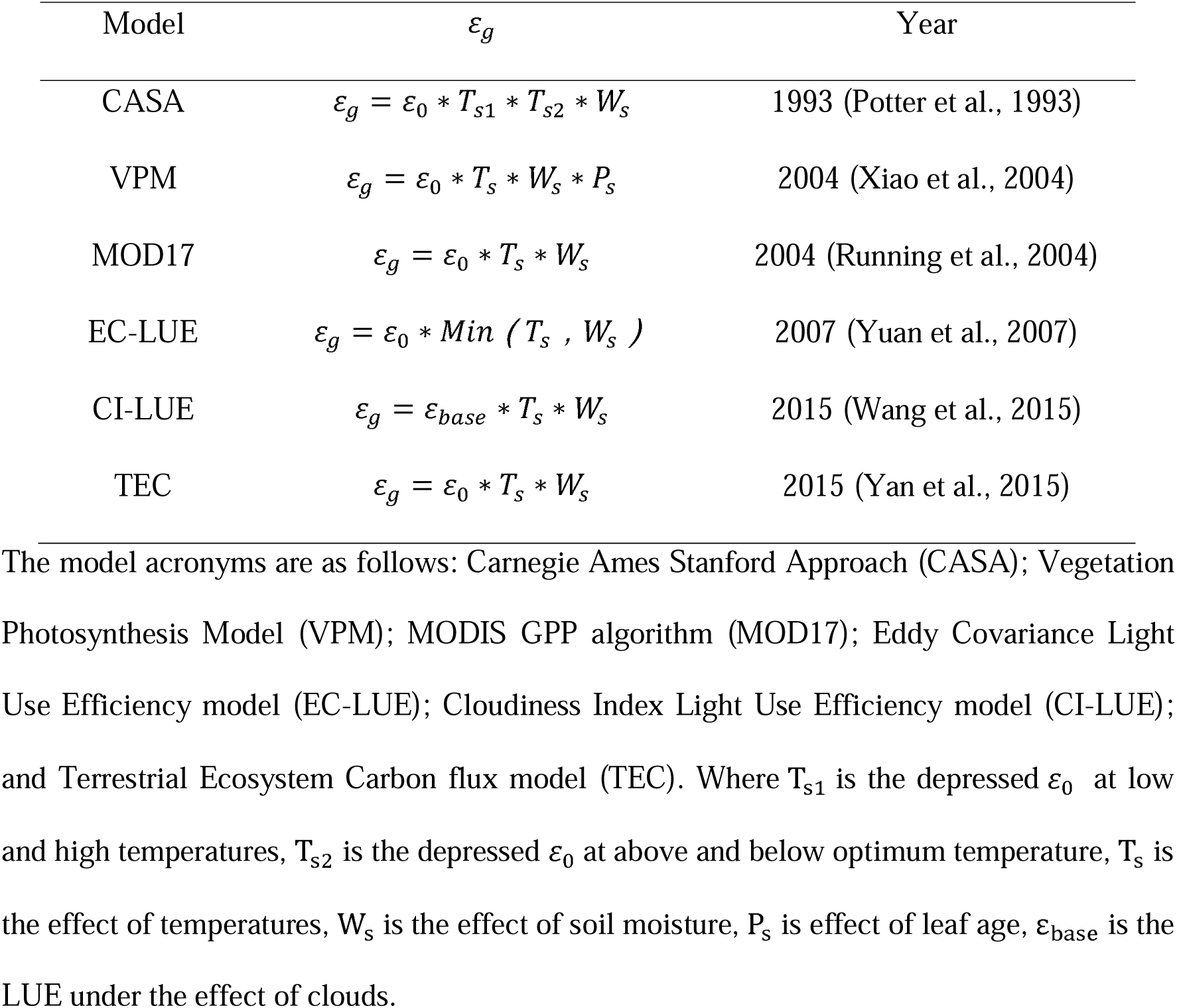
Common light use efficiency models.

| Model | $\varepsilon_g$ | Year |
| --- | --- | --- |
| CASA | $\varepsilon_g = \varepsilon_0 * T_{s1} * T_{s2} * W_s$ | 1993 (Potter et al., 1993) |
| VPM | $\varepsilon_g = \varepsilon_0 * T_s * W_s * P_s$ | 2004 (Xiao et al., 2004) |
| MOD17 | $\varepsilon_g = \varepsilon_0 * T_s * W_s$ | 2004 (Running et al., 2004) |
| EC-LUE | $\varepsilon_g = \varepsilon_0 * \text{Min} (T_s, W_s)$ | 2007 (Yuan et al., 2007) |
| CI-LUE | $\varepsilon_g = \varepsilon_{base} * T_s * W_s$ | 2015 (Wang et al., 2015) |
| TEC | $\varepsilon_g = \varepsilon_0 * T_s * W_s$ | 2015 (Yan et al., 2015) |

The model acronyms are as follows: Carnegie Ames Stanford Approach (CASA); Vegetation Photosynthesis Model (VPM); MODIS GPP algorithm (MOD17); Eddy Covariance Light Use Efficiency model (EC-LUE); Cloudiness Index Light Use Efficiency model (CI-LUE); and Terrestrial Ecosystem Carbon flux model (TEC). Where $T_{s1}$ is the depressed $\varepsilon_0$ at low and high temperatures, $T_{s2}$ is the depressed $\varepsilon_0$ at above and below optimum temperature, $T_s$ is the effect of temperatures, $W_s$ is the effect of soil moisture, $P_s$ is effect of leaf age, $\varepsilon_{base}$ is the LUE under the effect of clouds.

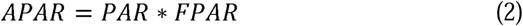

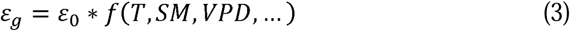

Where GPP is gross primary productivity, APAR is absorbed photosynthetically active radiation, PAR is photosynthetically active radiation, FPAR is the fraction of photosynthetically active radiation absorbed by vegetation, *ε_g_* is LUE under specific climatic conditions, *ε*_0_ is maximum LUE, T is temperature, SM is soil moisture, VPD is vapor pressure deficit, and … is other possible regulatory factors (such as radiation, nitrogen, and CO□, etc.).

Plant traits are physio-morpho-phenological qualities that indirectly impact fitness and performance by shaping the life history strategies of plants (Tyagi and Kumar, 2024). Plant traits can maximize ecosystem multifunctionality (Gross et al., 2017), and are a key framework for understanding ecosystem responses to global changes (Rao et al., 2024). Plant traits include whole-plant traits (e.g., life history, plant height), leaf traits (e.g., leaf dry-matter content, thickness), stem traits, below-ground traits, and regenerative traits (Pérez-Harguindeguy et al., 2013). These traits parameters control the growth (Falster et al., 2018), photosynthesis (Luo et al., 2021) and carbon sequestration capacity (Gillis et al., 2023; Yan et al., 2023) of vegetation, and therefore, the variability of LUE is closely related to them. In the study of mature Norway spruce, whose forests age ranges from 41 to 128, it was found that LUE increases with the increase in forests canopy height and age (Gspaltl et al., 2013). The same applies to deciduous forests, while in evergreen forests, LUE rose till reaching a peak at approximately 90 years and then gradually decreased with age (Xu et al., 2020). High tree species richness increases the vertical structural complexity of the canopy. High tree species richness maximizes light interception, and promoted a more stable spatiotemporal (encompassing both horizontal and vertical dimensions) distribution of light interception in tropical forests (Duarte et al., 2021). These ultimately reflect the increase in LUE. The growth in forest canopy height and age, along with the enhancement of tree species richness, will all lead to an increase in leaf area. Therefore, studies in boreal coniferous forests have indicated that LUE showed a strong nonlinear dependency on the leaf area (Launiainen et al., 2016). Solar-induced chlorophyll fluorescence (SIF) is the light that is re-emitted from the light energy absorbed by plants during photosynthesis, after excluding the portions that are used for photosynthesis and dissipated in the form of heat. SIF and LUE exhibit different correlations at the mid-day, daily, and monthly timescales (Lin et al., 2019). In temperate and boreal ecosystems, nitrogen controls canopy LUE and explain 71% of the variance (Kergoat et al., 2008). Therefore, plant traits are essential for the understanding of LUE.

Climatic conditions affect the expression of LUE, and this influence seems to be more complex than that of plant traits. Irrigation can lead to an increase in soil moisture, but this does not increase LUE in a loblolly pine plantation (Campoe et al., 2013). Deep research shows that LUE was most responsive to plant moisture indicators, least responsive to soil moisture (Zhang et al., 2015). In the study that analyzed the FLUXNET2015 data using the generalized linear mixed-effects model, it was found that the daytime temperature was proven to be the most significant predictor of LUE (Bloomfield et al., 2022). The impact of temperature on LUE is concentrated in spring, while in summer, LUE is generally driven by water stress in summer for forests and grasslands (Chen et al., 2024). Furthermore, plant traits are closely related to the climatic parameters (Cheng et al., 2022; Heilmeier, 2019), thus LUE is influenced by both. Scientifically understanding LUE is crucial for elucidating global carbon sequestration potential.

In this research, we analyzed the global patterns of forests LUE, and explored their main drivers. Using remote sensing data, we calculated the LUE of global forests and compared performance across five forest types. Finally, we applied generalized linear models (GLM) to identify the main drivers of LUE across spatial and temporal variations.

## 2. Materials and Methods

### 2.1. Data

#### 2.1.1. Datasets used to calculate LUE

Gross Primary Productivity (GPP, kg C m^-2^ 8d^-1^) and Fraction of Photosynthetically Active Radiation (FPAR, dimensionless unit) were respectively obtained from 8-day composite datasets MOD17A2H (Running, 2021) and MOD15A2H (Myneni, 2021), each with 500 meter pixel size. Due to data gaps (GPP: 2001-06-26, and 2022-10-16; FPAR: 2001-02-18, 2001-06-26, 2016-02-18, and 2022-10-16), linear interpolation was applied to fill in missing values. Each calendar year includes 46 images. To obtain daily GPP of 8-day, it is necessary to divide by 8 for the first 45 images and by 5 (6 in a leap year) for the final image.

Photosynthetically Active Radiation (PAR, MJ m^-2^) was estimated from incident shortwave radiation (SW, W m^-2^) as PAR = SW * 0.45. SW was obtained from an hourly time-averaged data collection (variable surface net downward shortwave flux) in Modern-Era Retrospective analysis for Research and Applications version 2 (MERRA-2) ((GMAO), 2015). The spatial resolution is 0.5 ° x 0.625 °. The dataset was integrated to match the GPP.

#### 2.1.2. Plant Traits datasets

Forests canopy height (HEIGHT, meter) data was obtained from the global canopy height map at 10□meter pixel size for the year 2020 (Lang et al., 2023). Forests age (AGE, years) was sourced from the global forests age dataset with a 1000 meter pixel size for the year 2010 (Besnard et al., 2021). Tree species richness (RICHNESS, dimensionless unit) was acquired from the global tree species richness map at 0.025 ° x 0.025 ° resolution for the year 2015 (Liang et al., 2022). RICHNESS is the diversity of tree species within a given area, rather than a trait of individual plants. Strictly speaking, RICHNESS is considered a community functional trait, while we grouped it together with other plant traits for the convenience of discussion. FPAR and Leaf Area Index (LAI, m^-2^ m^-2^) were taken from the MOD15A2H, using the mean values for 2022. Solar-induced chlorophyll fluorescence (SIF, W m^−2^ μm^−1^ sr^−1^) was retrieved from the global ‘OCO-2’ SIF data set (GOSIF) at 0.05 ° x 0.05 ° resolution for the year 2022 (Li and Xiao, 2019).

Additionally, four leaf traits were included: leaf nitrogen content per leaf dry mass (LNC, mg g^-1^, reflects the photosynthetic capacity), leaf phosphorus content per leaf dry mass (LPC, mg g^-1^, reflects the energy transfer and growth), leaf area per leaf dry mass (SLA, mm^2^ mg^-1^, reflects light capture efficiency and growth strategy), and leaf dry mass per leaf fresh mass (LDMC, g g^-1^, reflects tissue density and water retention capacity). Although the global leaf traits represent properties at the leaf level, they are essential because they capture the physiological mechanisms underlying photosynthesis. These traits were sources from global trait maps at 3000 meter pixel size for the year (Moreno-Martinez et al., 2018).

#### 2.1.3. Climatic conditions datasets

PAR was obtained from MERRA-2, which is the same dataset as MOD17A2H algorithm. Elevation was obtained from AG100 at 100 meter pixel size (JPL, 2014). Precipitation (PRE, mm) was obtained from ERA5-Land monthly averaged dataset at 0.1 ° x 0.1 ° resolution (Muñoz Sabater, 2019). We chose not to continue using MERRA-2 for the reason that its relatively low resolution compared to ERA5. Daytime and Nighttime Land surface Temperature (LST-D, LST-N, □) were obtained from MOD21C3 (Hulley, 2021). It should be noted that considering that the air temperature in ERA5 is used to calculate LUE in MOD17A2H, the temperature here is LST instead of air temperature. This can avoid over-fitting in subsequent analyses.

#### 2.1.4. Land cover types dataset

Forest types were obtained from MCD12Q1, which is a land cover types dataset at 500 meter pixel size yearly (Friedl, 2019). The first land cover types (IGBP classification) were chosen. All datasets were resampled to a uniform spatial resolution of 10 km, to ensure consistency across analyses.

### 2.2. Method

#### 2.2.1. Identification of forest and forest types

Figure 1 illustrates the flow chart. A total of 22 images, spanning from 2001 to 2022, were selected and divided into two segments starting in 2011. The mode of pixel values for each 11-image segment represented land cover types for each period (previous period and latter period). If both segments indicated the same forest type, the pixel was classified as that forest type. This method effectively reduces the impact of potential land cover misclassification in specific years on the final results.

**Figure 1.**
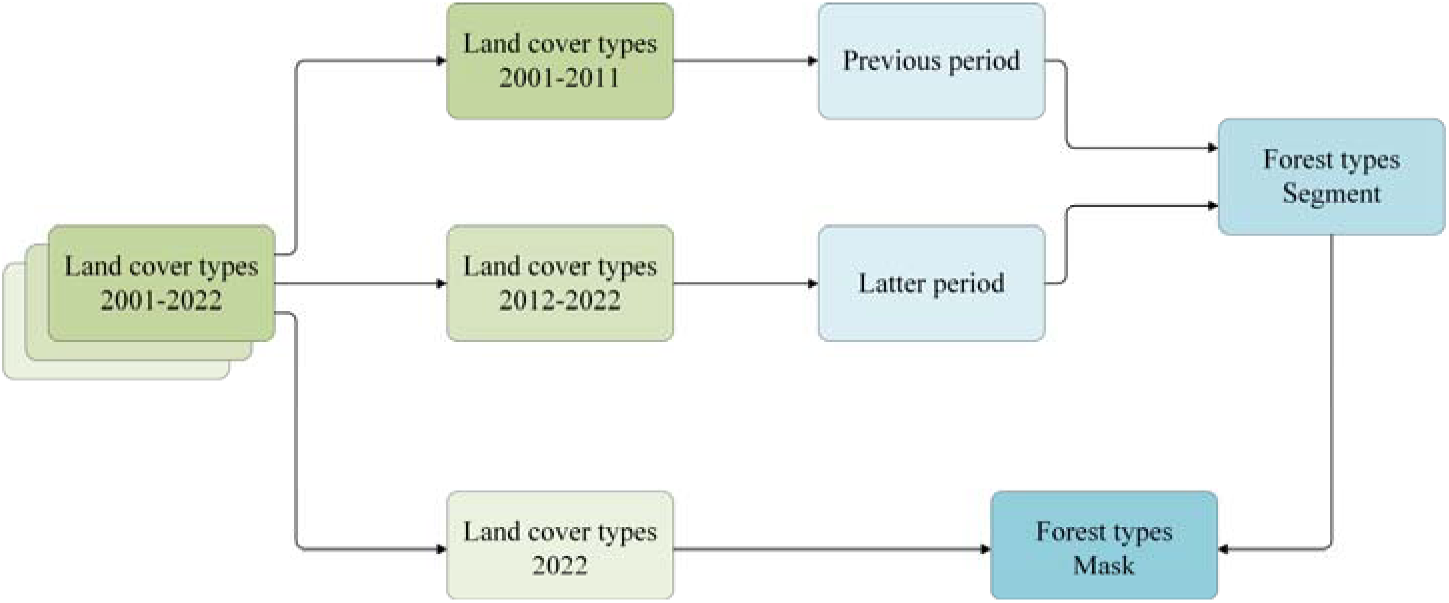
Identification of forest and forest types.

Moreover, forest loss in 2022 may have been overlooked in the previous analysis, so we applied a mask using 2022 forest type data to account for this.

#### 2.2.2. Calculation of LUE

LUE serves as an intermediate variable in the computation of GPP, although no direct data product exists for it. In this research, the calculation of LUE follows the logic in MOD17A2H product, in which land cover types, FPAR, PAR, Ts, and W_s_ are used as input variables to derive APAR and *ε_g_*, ultimately yielding GPP (Figure 2). Therefore, two approaches can be used to estimate LUE: (1) directly from environmental factors, as in the MOD17A2H algorithm, or (2) indirectly by derived it from GPP. We adopted the latter approach direct LUE observations for accuracy validation are unavailable, whereas GPP estimates are comparatively more reliable. This approach is therefore logically consistent.

**Figure 2.**
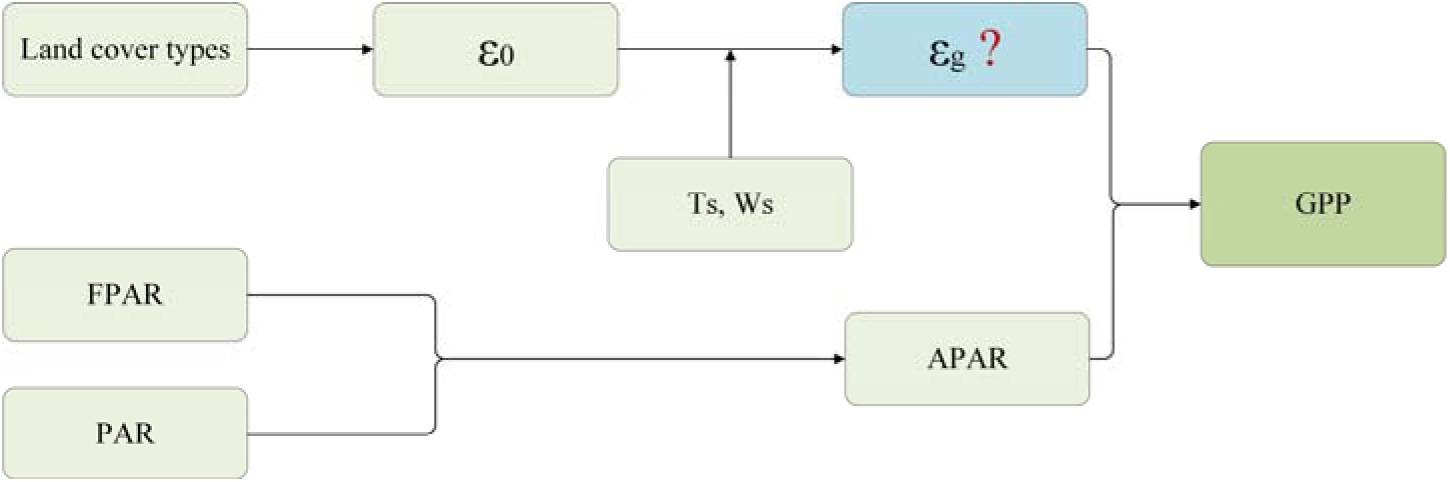
Algorithm theoretical basis of MOD17A2H. There is no data product about *ε_g_*.

The MOD12A2H GPP dataset has undergone extensive validation, demonstrating satisfactory performance (Zhang et al., 2017), without overall bias (Turner et al., 2006), and an ability to capture broad GPP trends (Tang et al., 2015) and seasonal variations (Sjöström et al., 2013) on an 8-day time scale. A critical evaluation of 45 global GPP products showed that, the estimates of the GPP products were uneven and different in vegetation types (Zhang and Ye, 2021), but we can find GPP generally performed well for forests from the figure. These findings suggest that the LUE values derived from GPP are reliable for forest ecosystems.

In the above logic, LUE is obtained by the following formula 4:

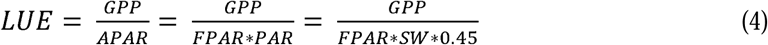

Note the units of the formula. LUE (g C MJ^-1^), GPP (g C m^-2^ d^-1^ from kg C m^-2^ 8d^-1^), FPAR (dimensionless unit), and PAR (SW, MJ m^-2^ d^-1^ from W m^-2^).

The calculation of LUE is done on Google Earth Engine (https://earthengine.google.com/).

#### 2.2.3. Analysis of distribution and influencing factors of LUE

We assessed the distribution of LUE, and analyzed its interannual variation using linear regression model. To smooth multi-year temporal variation curve of LUE, we used a moving average method with a window size of 4.

The research mainly examines the influence levels of different factors, such as plant traits and climatic conditions. All variables were normal-score normalized, and the dataset was split into 80% training data and 20% validation subsets. Generalized linear model (GLM) was employed to quantify the influence and interpretative contribution of each factor to LUE. LUE variation in forests was analyzed across both spatial and temporal dimensions. Spatial analysis includes all forest pixel, and focused on the 2022 annual LUE and incorporated all plant traits and climatic conditions. While temporal analysis includes every eight days data from 2001 to 2022, and considered a subset of continuous variables—FPAR, LAI, PAR, PRE, LST-D, and LST-N.

## 3. Results

### 3.1. Distribution of global forests and forest types

The distribution of global forests is uneven, with major concentrations in Northern South America, Central Africa, Southeast Asia, Northern Russia and Canada, and the Pacific and Atlantic coasts (Figure 3 a). Evergreen broadleaf forests (EBF) dominate the equatorial region, comprising 46.63% of global forest area. In contrast, evergreen needleleaf forests (ENF), deciduous needleleaf forests (DNF), and mixed forests (MF) are prevalent in the mid- and high-latitude regions of the Northern Hemisphere, collectively known as boreal forests. Positioned between tropical and boreal forests are temperate forests, primarily composed of deciduous broadleaf forests (DBF).

**Figure 3.**
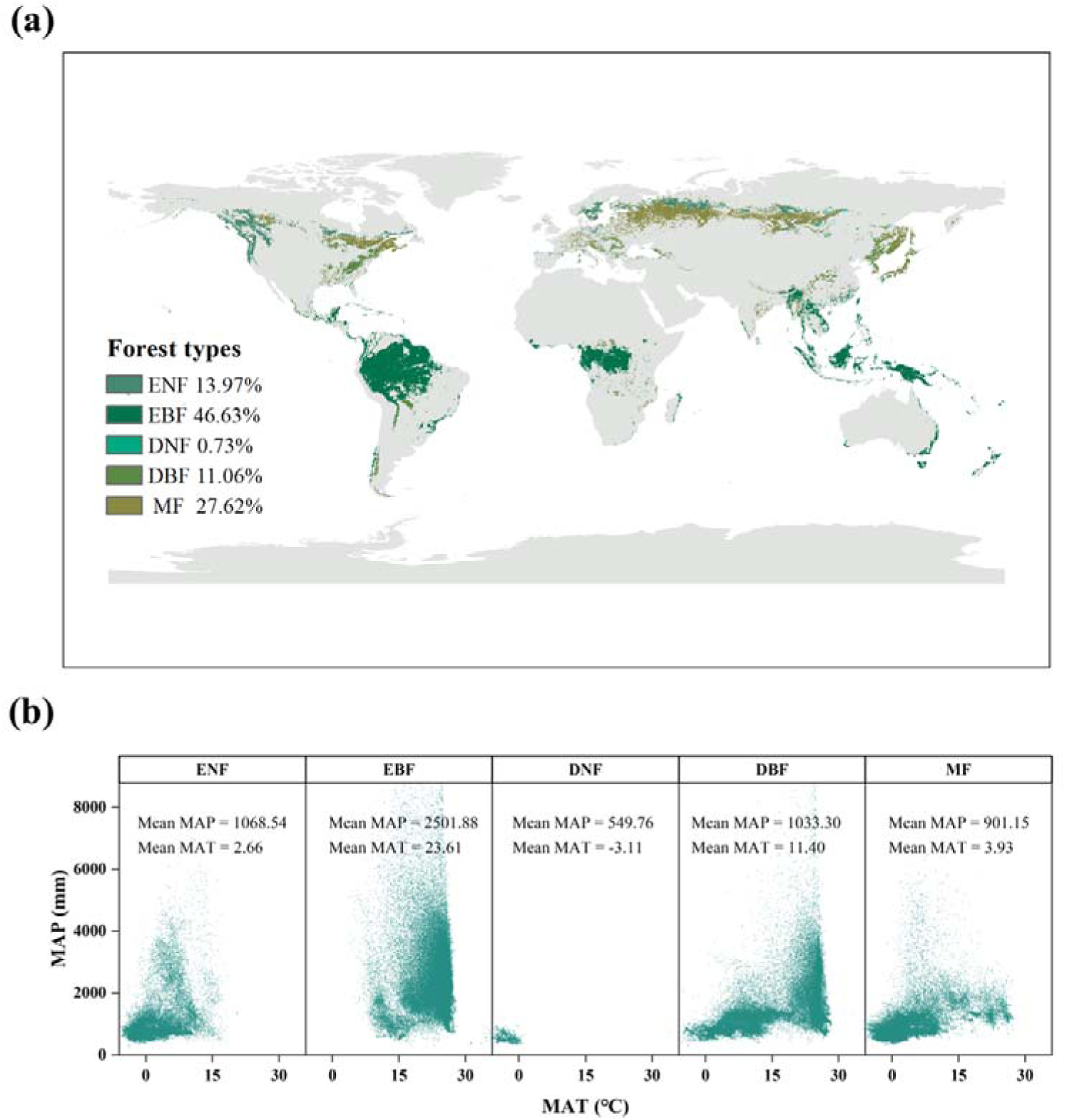
Global distribution of forests and forest types (a) and meteorological conditions in the habits of different forest types (b). The spatial resolution of the pixels/points is 10 km. ENF, evergreen needleleaf forests; EBF, evergreen broadleaf forests; DNF, deciduous needleleaf forests; DBF, deciduous broadleaf forests; MF, mixed forests. MAP, mean annual precipitation (mm); MAT, mean annual temperature (°C).

Different meteorological conditions breed different forest types (Figure 3 b). EBF thrive in high temperature and humidity regions, while the cold, dry climate of northern regions favors the development of coniferous forests.

### 3.2. Spatial and temporal variations of LUE

LUE gradually decreased from equator to the poles, ranging from 0 to 1.55 g C MJ^-1^ (Figure 4 a). LUE levels vary among forest types (Figure 4b), with EBF exhibiting the highest LUE (1.26 g C MJ^-1^), followed by ENF at 0.76 g C MJ^-1^. DBF and MF show similar levels at 0.50 g C MJ^-1^, while DNF have the lowest LUE at 0.41 g C MJ^-1^.

**Figure 4.**
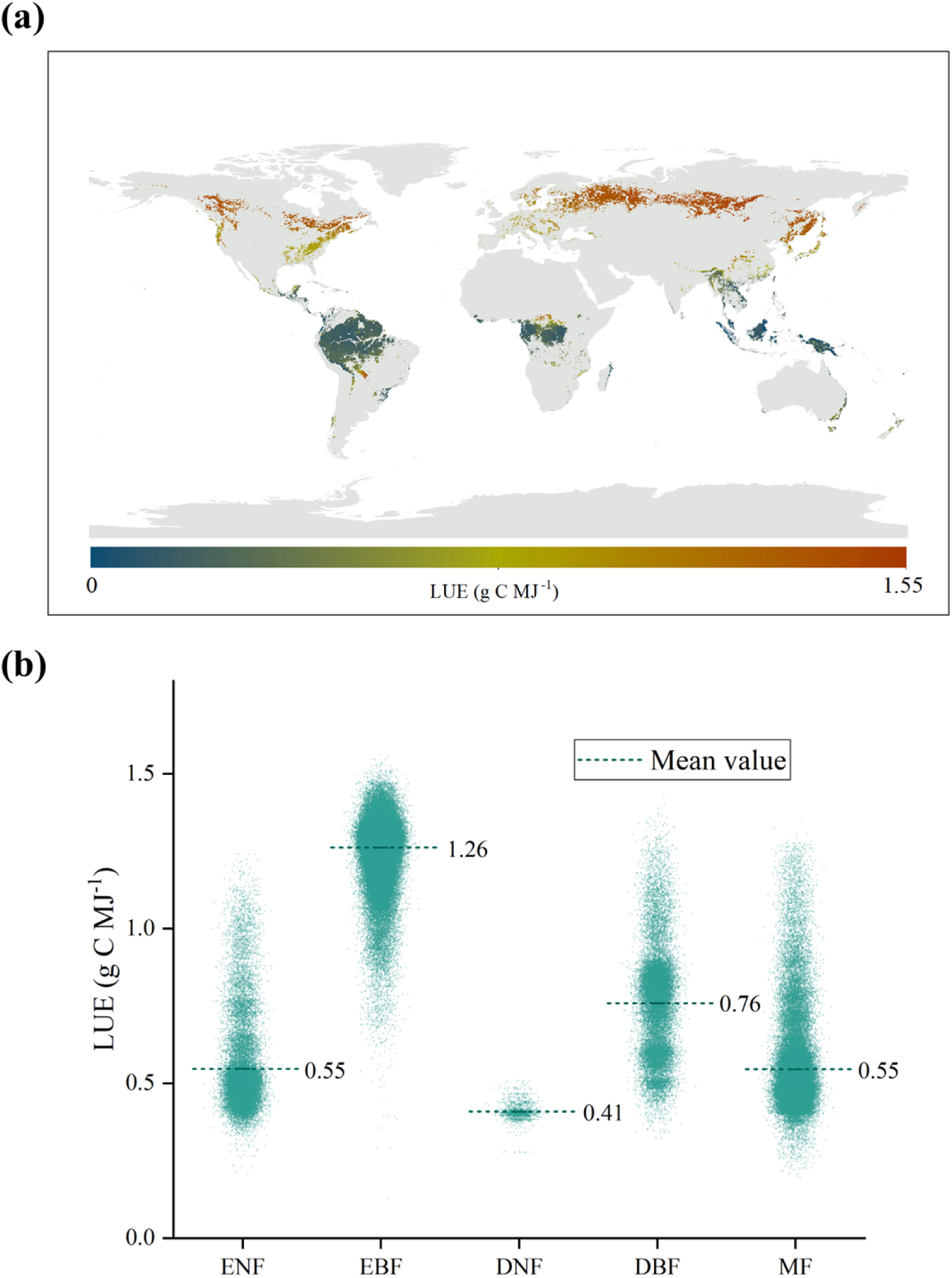
Distribution of LUE in forests (a) and LUE value of different forest types (b) during the period 2001-2022. ENF, evergreen needleleaf forests; EBF, evergreen broadleaf forests; DNF, deciduous needleleaf forests; DBF, deciduous broadleaf forests; MF, mixed forests.

The global average annual LUE of forests is 0.93 ± 0.36 g C MJ^-1^ during the period 2001-2022. Interannual LUE show a significant increase trend, with a trend of 0.0034 g C MJ^-1^ yr^-1^ (Figure 5).

**Figure 5.**
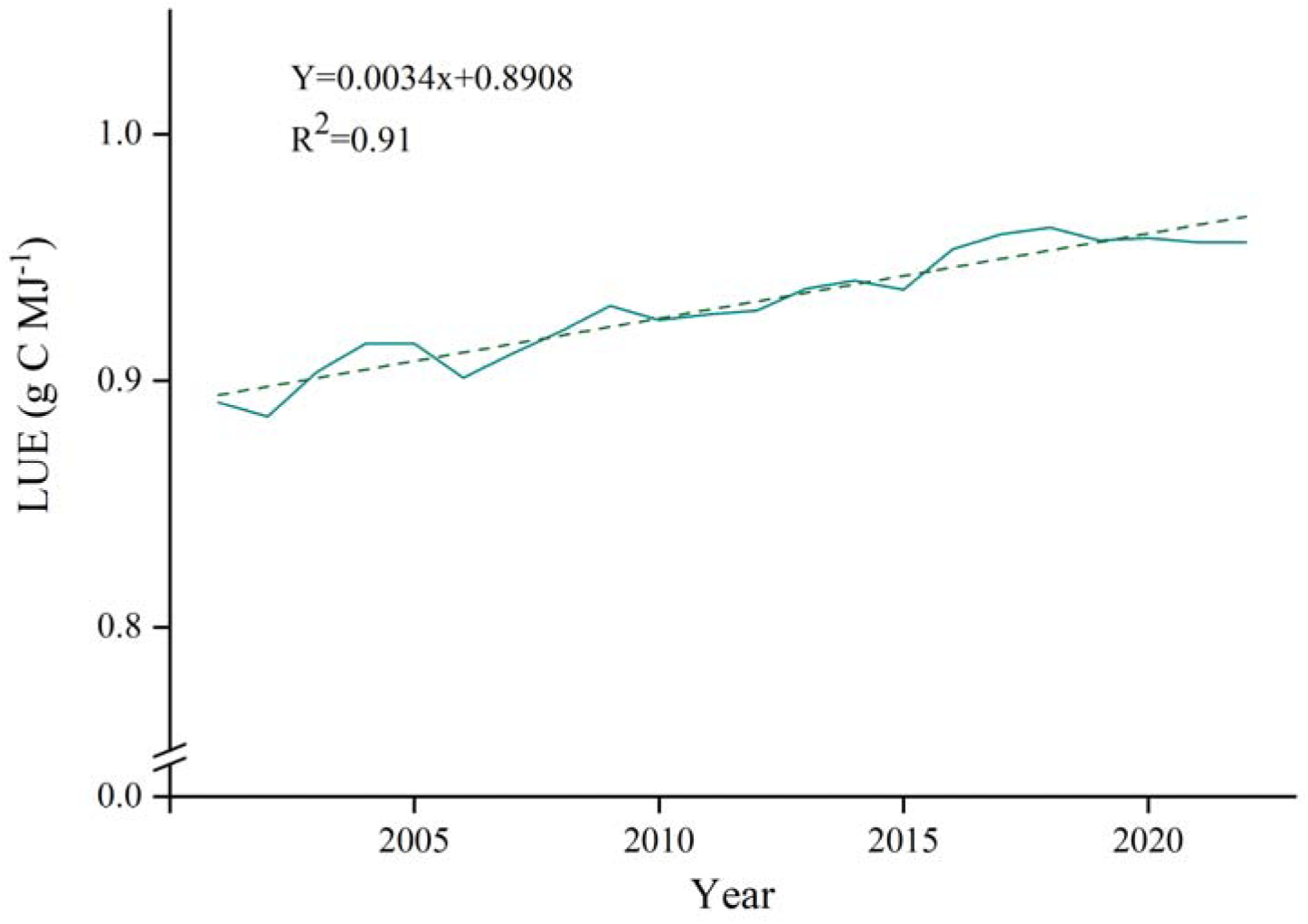
Temporal variation trend of LUE in forests (annual scale). The dotted line is the annual curve of linear fitting.

8-day LUE in forests is presented in two segments from 2011. The superimposed curves of these two segments show that the LUE in the latter 11 years is higher than that in the previous 11 years, both low and high values (Figure 6).

**Figure 6.**
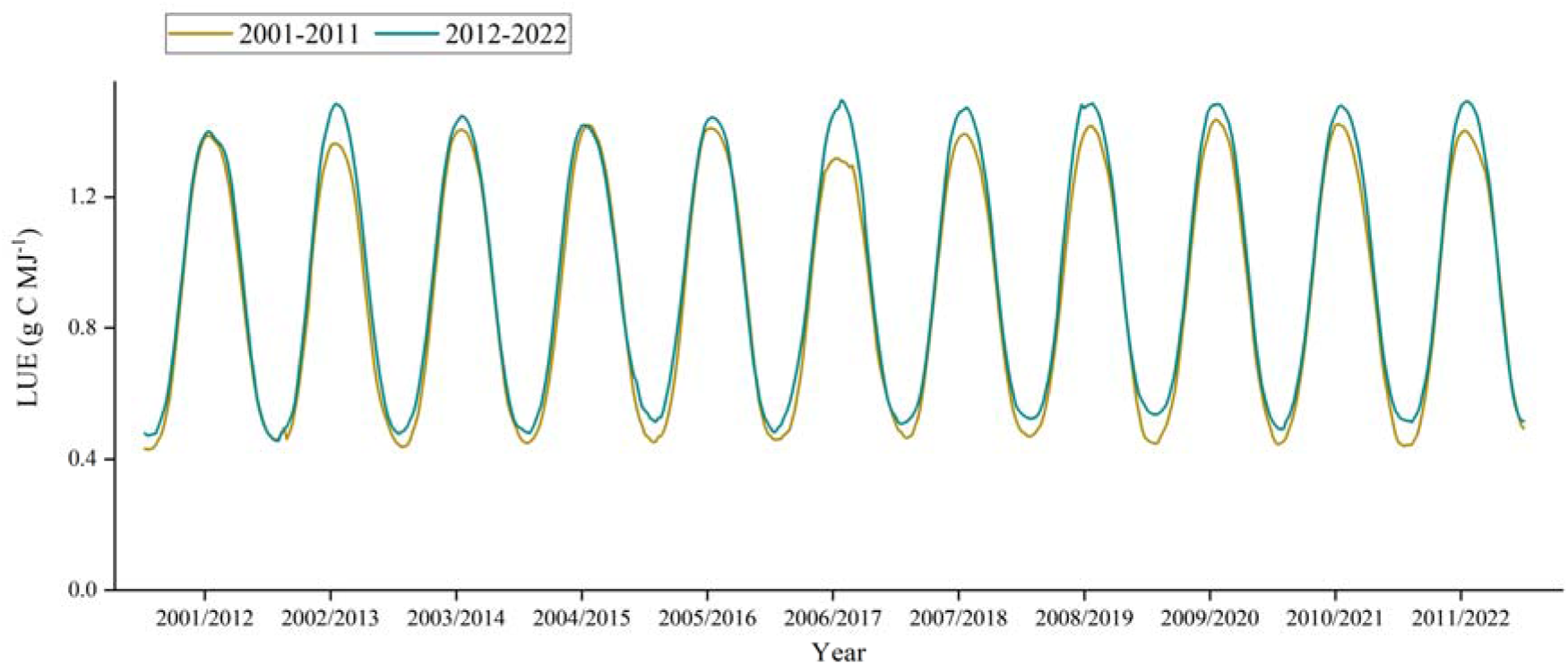
Temporal variation trend of LUE in forests (8-day scale).

Temporal variation trend of LUE across forest types aligns with the spatial distribution patterns (Figure 7 a). In descending order, they are EBF, ENF, DBF, MF, and DNF. EBF shows minimal seasonal variation in LUE, while the other four forest types display distinct seasonal dynamics. The LUE magnitudes in ENF, MF, and DNF are similar, all following unimodal seasonal curves. Notably, DNF’s LUE approaches zero in the Northern Hemisphere winter, while DBF exhibits a non-unimodal curve, likely due to its broad distribution and phenological diversity.

**Figure 7.**
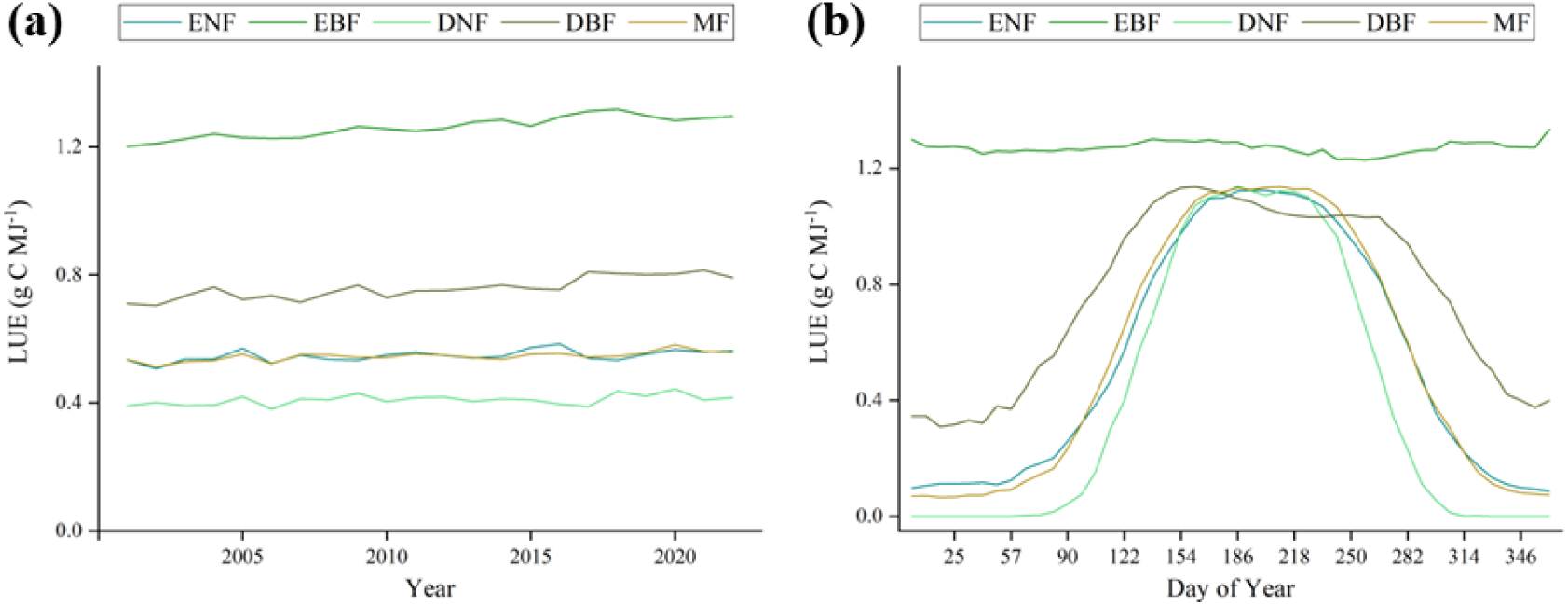
Temporal variation trend of LUE of different forest types. (a) annual scale, (b) 8-day scale average across 2001-2022.

### 3.3. Analysis of influencing factors of LUE

#### 3.3.1. Spatial analysis

Spatial analysis reveals correlations between LUE, plant traits, and climatic conditions (Figure 8). Variables strongly correlated with LUE include LST-D, LST-N, LAI, PAR, RICHNESS, FPAR, AGE, SIF, MAP, HEIGHT, and LNC (Figure S1). Conversely, LPC, SLA, ELE, and LSMC exhibit weaker correlations with LUE. Additionally, some variables are strongly intercorrelated, such as FPAR and LAI, and LST-D and LST-N.

**Figure 8.**
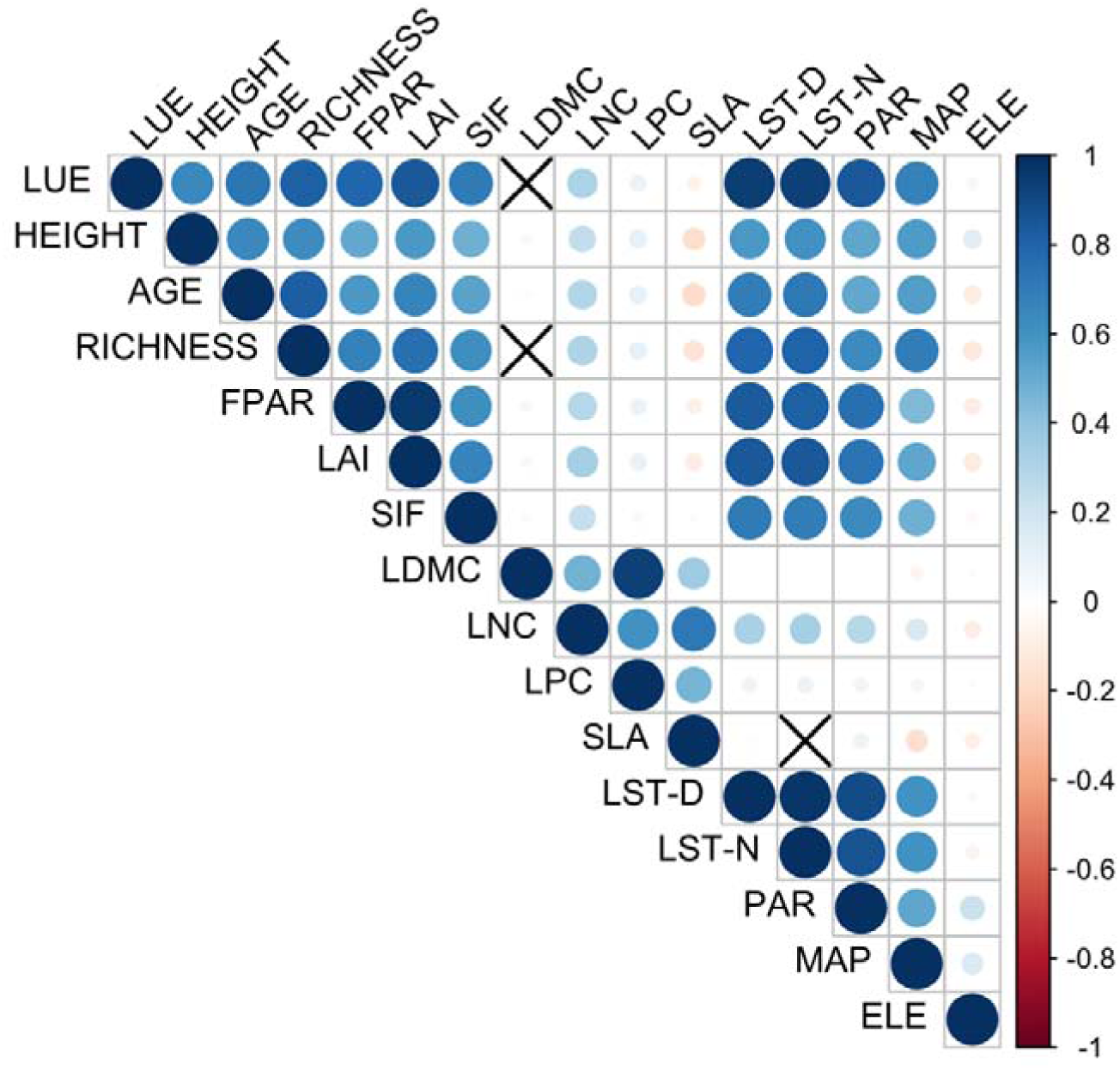
Correlation matrix plot of LUE and multiple plant traits and climatic conditions. × means not significant.

There are slight differences among different forest types (Figure S2). EBF and DNF exhibited lower correlations to LUE compared with the other three forest types. PAR is a negative factor to LUE in EBF. In addition, AGE also has a non-positive effect to LUE for all forest types except EBF.

Both plant traits and climatic conditions influence the spatial variation of LUE in forests (Table 2). LAI dominated the spatial variation of LUE. HEIGHT, AGE, RICHNESS, LDMC, LNC, and LST-D play a role to some extent. FPAR, LPC, MAP, and SIF play a negative role in the spatial variation of LUE. The built generalized linear model accounted for 86% of the spatial variation of LUE (Figure 9).

**Figure 9.**
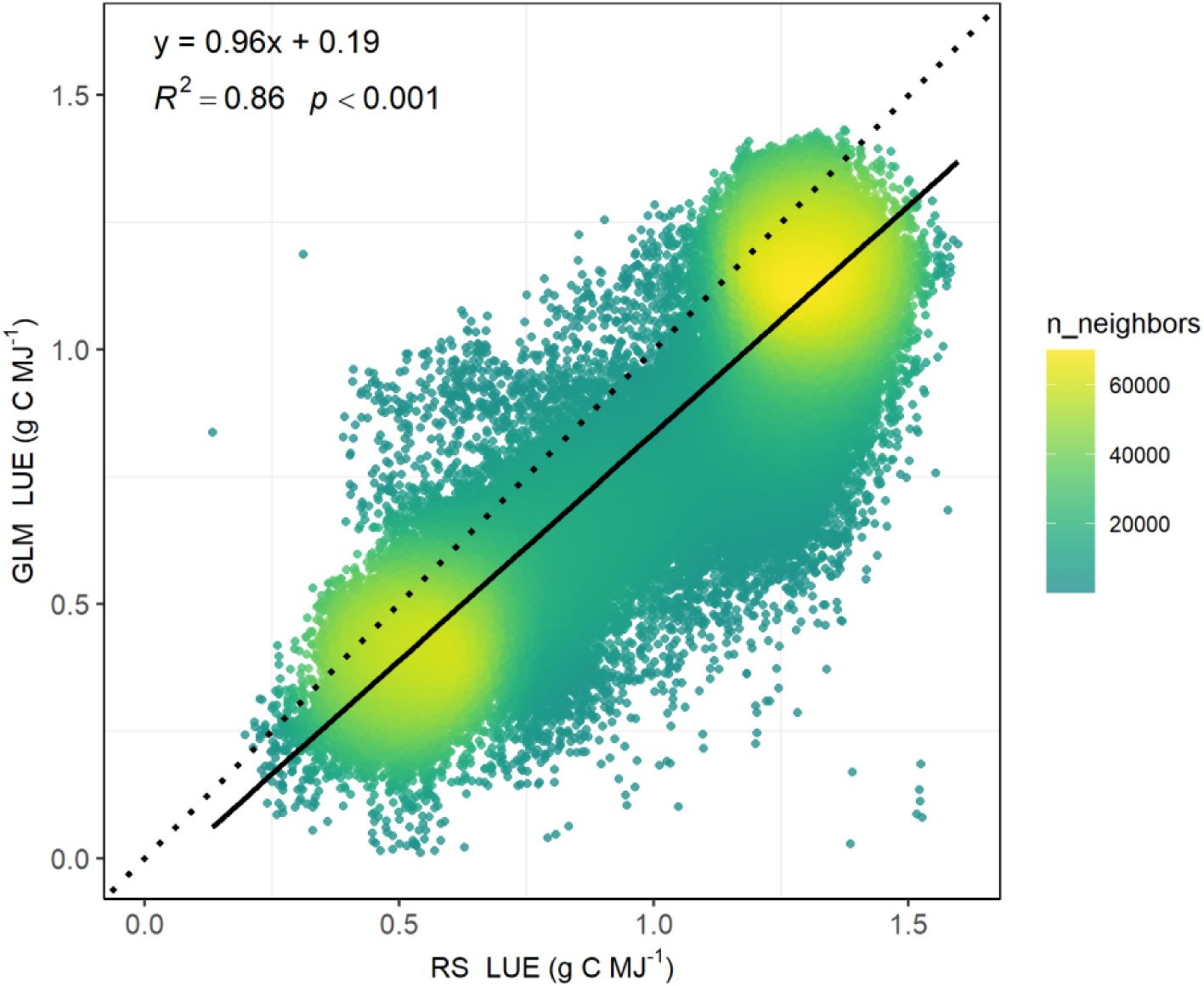
Scatter plot of RS (remote sensing calculated) LUE and GLM (generalized linear model calculated) LUE.

**Table 2.** Drivers and standardized coefficient of LUE by a generalized linear model. × means not significant.

| Plant Traits | HEIGHT | AGE | RICHNESS | FPAR | LAI | SIF | LDMC | LNC | LPC | SLA |
| --- | --- | --- | --- | --- | --- | --- | --- | --- | --- | --- |
| Standardized Coefficient | 0.13 | 0.09 | 0.16 | -0.30 | 0.42 | -0.04 | 0.14 | 0.09 | -0.17 | 0.01 × |
| Climatic Conditions | LST-D | LST-N | PAR | MAP | ELE |  |  |  |  |  |
| Standardized Coefficient | 0.18 | 0.10 | 0.05 | -0.06 | 0.05 |  |  |  |  |  |

#### 3.3.2. Temporal analysis

There is a correlation among LUE, FPAR, LAI, PAR, PRE, LST-D, and LST-N in temporal analysis (Figure 10). Each variable is associated with temporal variation of LUE (Figure S3). In addition, these variables are clearly correlated with each other, such as FPAR, LAI, and PAR.

**Figure 10.**
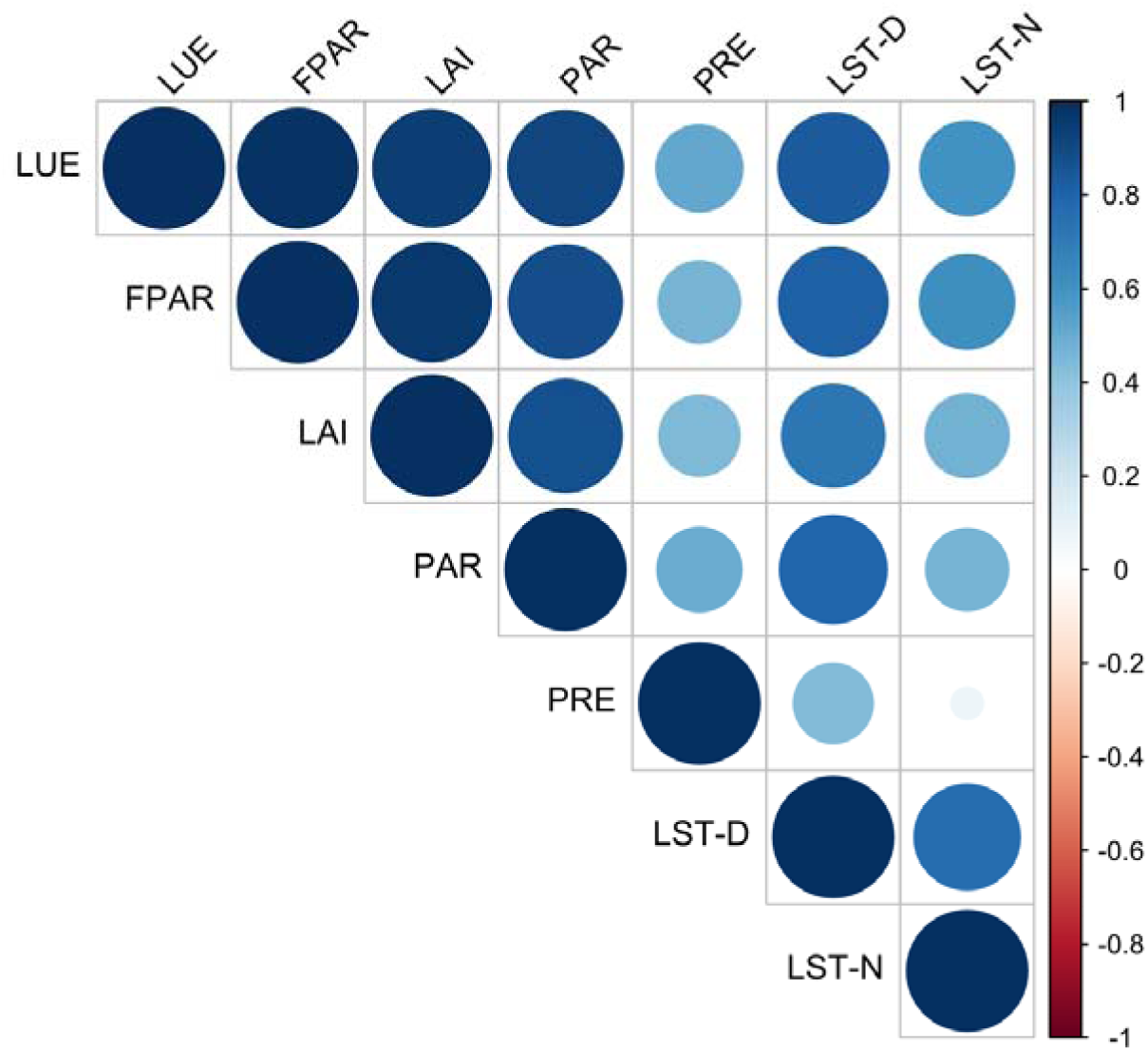
Correlation matrix plot of LUE and multiple plant traits and climatic conditions. × means not significant.

The patterns are consistent across different forest types, except for EBF (Figure S4). In EBF, PAR makes a strong negative contribution to LUE, while the positive effects of PRE and LST-D are also lower than those in other forest types.

Several variables mentioned above influence the temporal variation of LUE in forests (Table 3). FPAR dominated the temporal variation of LUE. PAR and PRE play a role to some extent. LAI play a negative role in the spatial variation of LUE. LST has no significant effect. The built generalized linear model accounted for 98% of the temporal variation of LUE (Figure 11).

**Figure 11.**
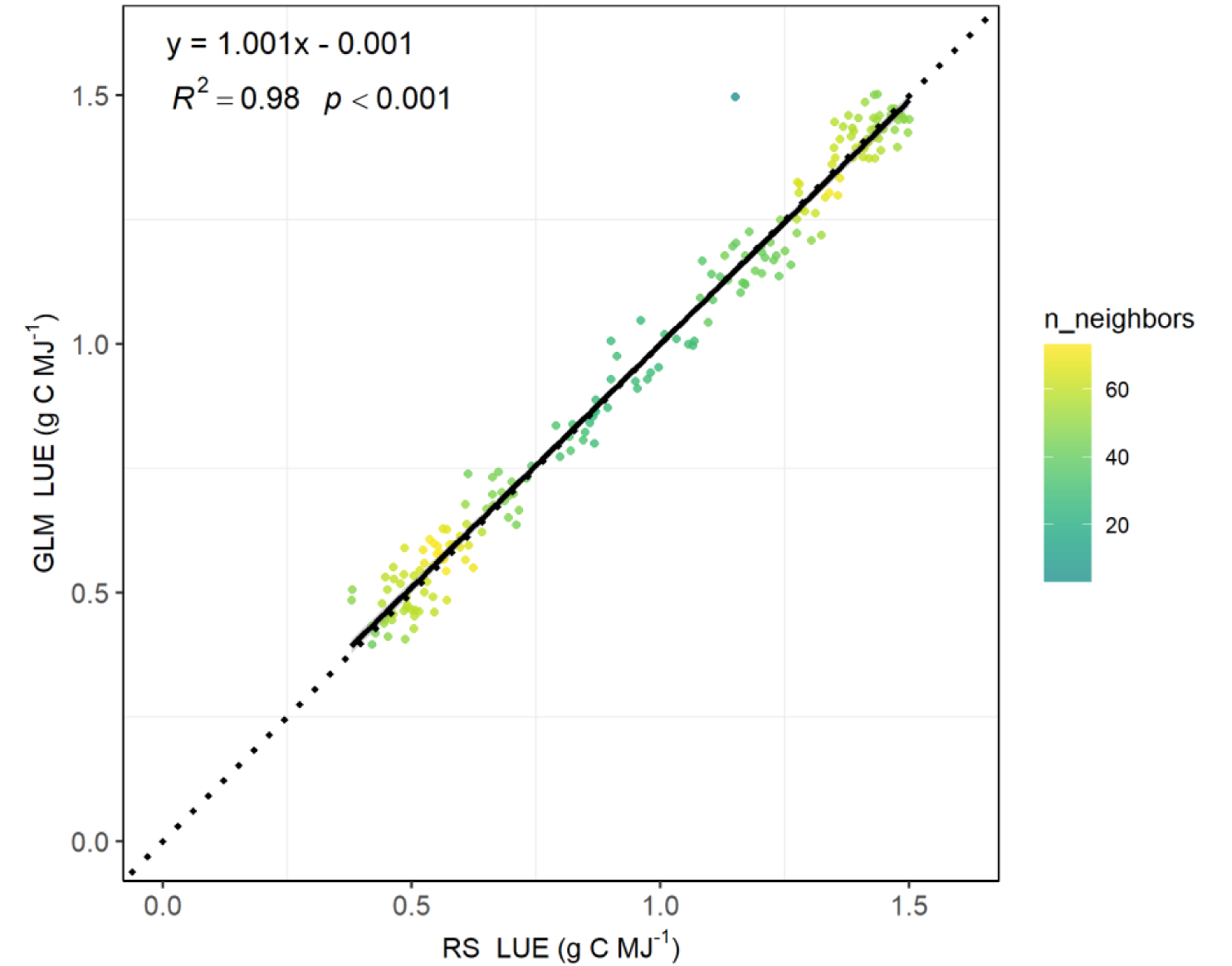
Scatter plot of RS (remote sensing calculated) LUE and GLM (generalized linear model calculated) LUE.

**Table 3.** Drivers and standardized coefficient of LUE by a generalized linear model. × means not significant.

| Climatic Conditions | FPAR | LAI | PAR | PRE | LST-D | LST-N |
| --- | --- | --- | --- | --- | --- | --- |
| Standardized Coefficient | 0.91 | -0.14 | 0.18 | 0.06 | 0.02× | 0× |

## 4. Discussion

In recent years, LUE has gained significant attention in research. It serves as a critical variable to evaluate the physiological effects of elevated CO_2_ on land–atmosphere carbon and water exchange (Zhan et al., 2022) and help explain the mechanism behind the pattern that the increase of global GPP is higher than earth greening in drylands (Wang et al., 2023). Examining LUE variation patterns enhances our understanding and ability to predict climate change impacts on ecosystem carbon and water cycles.

### Variations of LUE in forests

The global average annual LUE of forests is 0.93 ± 0.36 g C MJ^-1^ during the period 2001-2022. This estimate is within the range of previous research (about 0.4 to 1.2 g C MJ^-1^) with data-driven models and process-based models for 1982–2013 (Tang et al., 2020). This value is lower than LUE (1.36 ± 0.46 g C MJ^-1^) in four forest types based on flux-tower sites for 2000-2014 (He et al., 2022). The flux tower observation usually conducted on representative vegetation, and there is basically no forest loss around the towers. This means that the measurements are biased toward relatively undisturbed or mature forests, which may have higher LUE. At the same time, the research mentioned above focus on LUE during the growing season, and do not include DNF whose LUE is lowest in our research. These two points are probably the main reason for the low value in our study. Temporal variation trend of LUE in our research (Figure 7 b) is close to that on a research about monthly dynamic variation (Wei et al., 2017). Both researches show that LUE in middle to high latitudes forests (ENF, DBF, and MF) display an obvious seasonal change, while no apparent temporal trend of LUE is found in low latitude forests (EBF). The comparison results show that the LUE calculated in this research in reliable. Considering global forest pixels, this research can fully reflect the variation in LUE of forest ecosystem.

LUE of forests increases with a trend of 0.0034 g C MJ^-1^ yr^-1^. Satellite-derived datasets also show widespread increasing LUE of terrestrial ecosystem (about 0.0012 g C MJ^-1^ yr^-1^) over the past two decades, which is driven by nitrogen deposition and CO_2_ fertilization effects, especially in evergreen broadleaf forests (0.0060 g C MJ^-1^ yr^-1^) (Liu et al., 2024). On a longer time scale (from 1982 to 2018), studies on grassland ecosystems in Northern China have also demonstrated a substantial rising trend for LUE (0.0034 g C MJ^-1^ yr^-1^) (Yuan et al., 2023). Maximum LUE has seasonal fluctuations (Lin et al., 2017), and maximum LUE of shaded leaves 2 to 4 times that of sunlit leaves (Zhou et al., 2016). Both have important influence on the estimation of GPP. A fully understanding of variation of LUE will help to improve the measurement of ecosystem carbon sinks.

### LUE is jointly driven by Plant traits and climatic conditions

LUE is generally significantly controlled by climate (radiation, temperature and precipitation). Globally, higher precipitation correlates with increased LUE, and higher temperatures are associated with elevated LUE primarily in the mid to high latitudes of the Northern Hemisphere (Tang et al., 2020). However, the spatial correlation between LUE and precipitation in forests differs from previous studies; mean annual precipitation (MAP) shows a negative correlation with LUE spatial variation. This may be because precipitation is not a limiting factor in favorable forest climates. Conversely, precipitation plays a more prominent role in temporal LUE variation, relating closely to wet and dry seasons, as observed in four contrasting forest ecosystems in Southwest China (Fei et al., 2019). Spatially, LST has a stronger influence on LUE than it does temporally, where its effect is insignificant. Additionally, spatial LUE is more influenced by PAR.

Plant traits is also a vital influencing variable for LUE. However, relatively little research has been focused on the this. In this research, plant traits contribute a large proportion to the spatial variation of LUE. LAI is the main positive influencing variables of spatial variation of LUE in forests, even more than climatic conditions. This is different from the fundings that water factors is more important for multiple vegetation types (He et al., 2022). However, LAI is negative to LUE at temporal level. That may reflect that in forest ecosystems (higher LAI compared to other vegetation types), progressively higher LAI does not necessarily enhance LUE. RICHNESS also has a positive influence in spatial variation of LUE. Mixing tree species increases tree productivity, and enhance tree performance (Depauw et al., 2024). This enhancement may be achieved by increasing light interception and thereby improving LUE. FPAR is the main negative influencing variables for spatial variation of LUE, while it is opposite for temporal variation of LUE. FPAR changes with vegetation growth stage and season, which can reflect the temporal dynamics of LUE. It’s noteworthy that the relationship between leaf traits (LDMC, LNC, LPC, and SLA) and LUE is not strong. This phenomenon deserves attention. Leaves is the place of photosynthesis, and the element content in it should have a closer relationship with LUE. It can be seen from this that in the existing LUE calculations, leaf traits are seriously lacking. Therefore, taking into account plant traits holds positive implications for estimating GPP with LUE models, which are insufficiently considered in current models. Only a little of the situation of the vegetation itself is considered in the model, and the parameters tend to represent the biome-specific factors rather than species-specific efficiency factors (Ahl et al., 2004). This was already raised in 2004, while it has not been completely resolved even to this day. Given the increasingly extensive demand of high-resolution carbon mapping nowadays, this situation merits particular attention.

LUE is jointly driven by plant traits and climatic conditions. The GLM models established by them can well reflect the spatial and temporal variation of LUE. However, there are also some limitations in this research. For instance, most of the dataset are calculated from spectral reflectance data. Although each dataset has been verified, there is inevitably a certain degree of autocorrelation, which is also the limitation of remote sensing technology itself.

### LUE is one of optically distinguishable functional traits that can be quantified

Remote sensing facilitates the assessment of plant traits through optical principles, leading to the concept of optically distinguishable functional traits (Ustin and Gamon, 2010). However, classification based on plant traits does not fully capture variations in photosynthesis and transpiration (Groenendijk et al., 2011). Additionally, plant traits include numerous variables that are challenging to standardize. LUE is one possible composite indicator, which is driven by plant physiological and climatic conditions, and considering plant traits themselves. As an optically distinguishable variable, LUE can be derived from reflectance or vegetation indices (Zhang et al., 2018) without factoring in climatic conditions. In particular, the photochemical reflectance index (PRI) correlates strongly with LUE on short (daily) timescales in mixed (Ma et al., 2020), deciduous, and evergreen broadleaf forests (Soudani et al., 2014), though this relationship requires validation across multiple vegetation types and larger scales.

## 5. Conclusion

LUE is the fundamental parameter for GPP modeling. However, it has not received adequate attention. In this study, we conducted a comprehensive analysis of LUE in global five forest types, highlighting significant spatial and temporal variations influenced by plant traits and climatic conditions. The global average annual LUE of forests, estimated at 0.93 ± 0.36 g C MJ^-1^, demonstrates an increase trend from 2001 to 2022. EBF exhibited the highest LUE, whereas DBF had the lowest. The incorporation of multiple factors in GLM successfully explained a substantial portion of the variations of LUE in temporal and spatial patterns, respectively, emphasizing the synergistic contribution between biological and climatic drivers. Our findings underscore the necessity of integrating detailed plant traits and climatic variables to enhance the predictive ability of carbon cycle models. Future work should focus on exploring the potential impacts of climate change on LUE to further facilitate sustainable forest management and mitigate climate change

## Data availability statement

Data sets utilized for this research are as follows:

GPP dataset, https://doi.org/10.5067/MODIS/MOD17A2H.061;

FPAR and LAI dataset, https://doi.org/10.5067/MODIS/MOD15A2H.061;

SW dataset, https://doi.org/10.5067/Q9QMY5PBNV1T;

Forests canopy dataset, https://doi.org/10.3929/ethz-b-000609802;

Forests age, https://doi.org/10.17871/ForestAgeBGI.2021;

Tree species richness, https://doi.org/10.6084/m9.figshare.17232491.v2;

SIF dataset, https://data.globalecology.unh.edu/data/GOSIF_v2/;

Leaf traits, https://www.try-db.org/TryWeb/Data.php#59;

Elevation, https://doi.org/10.5067/COMMUNITY/ASTER_GED/AG100.003;

Precipitation, https://doi.org/10.24381/cds.68d2bb30;

Daytime and Nighttime Land surface Temperature, https://doi.org/10.5067/MODIS/MOD21C3.061;

Forests types, https://doi.org/10.5067/MODIS/MCD12Q1.061.

