## Appendix S1 for "Global variations of Light Use Efficiency in Forests Jointly Driven by Plant Traits and Climatic Conditions"

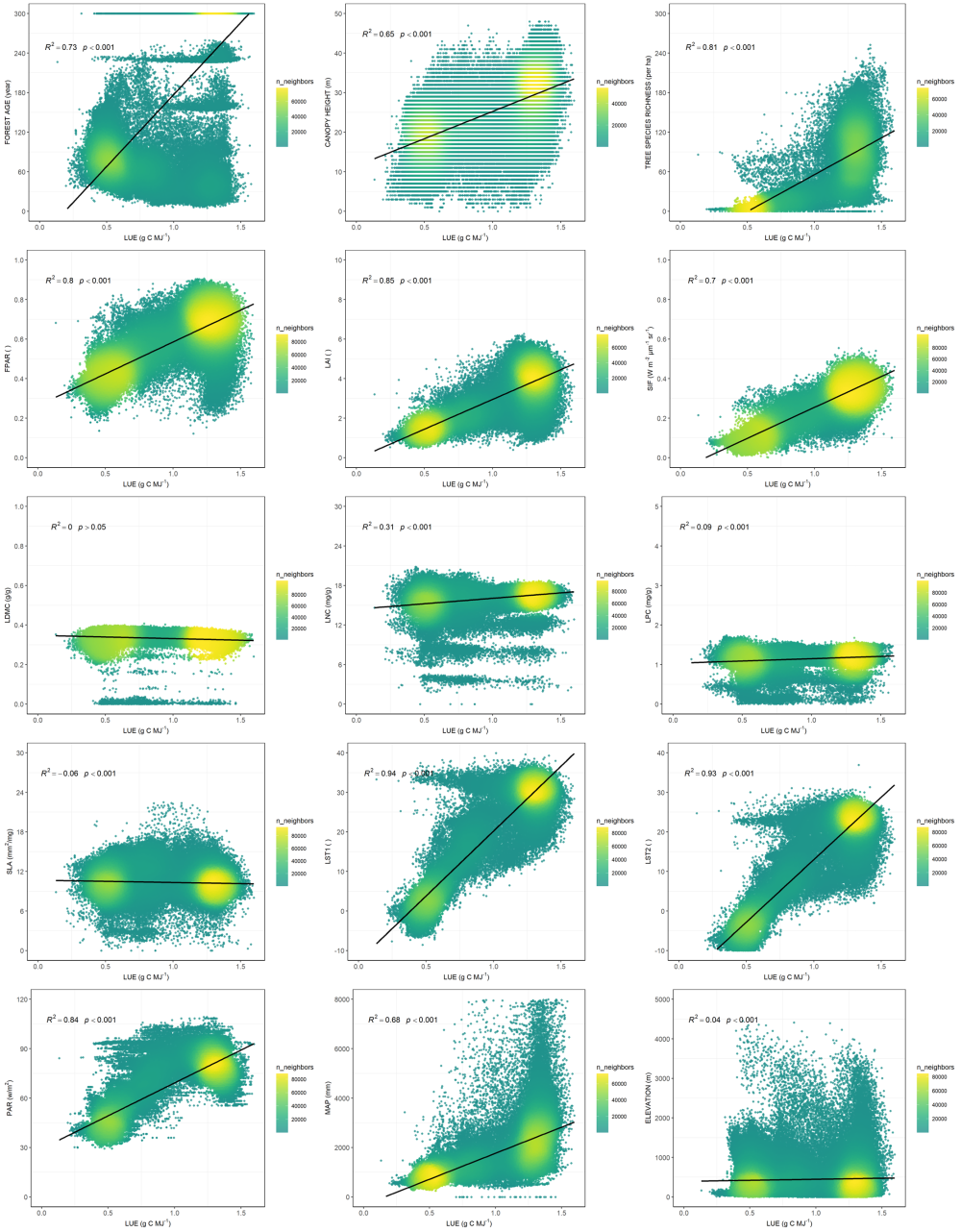

Figure S1 Scatter plot of LUE and multiple plant functional traits and environmental conditions in spatial analysis.

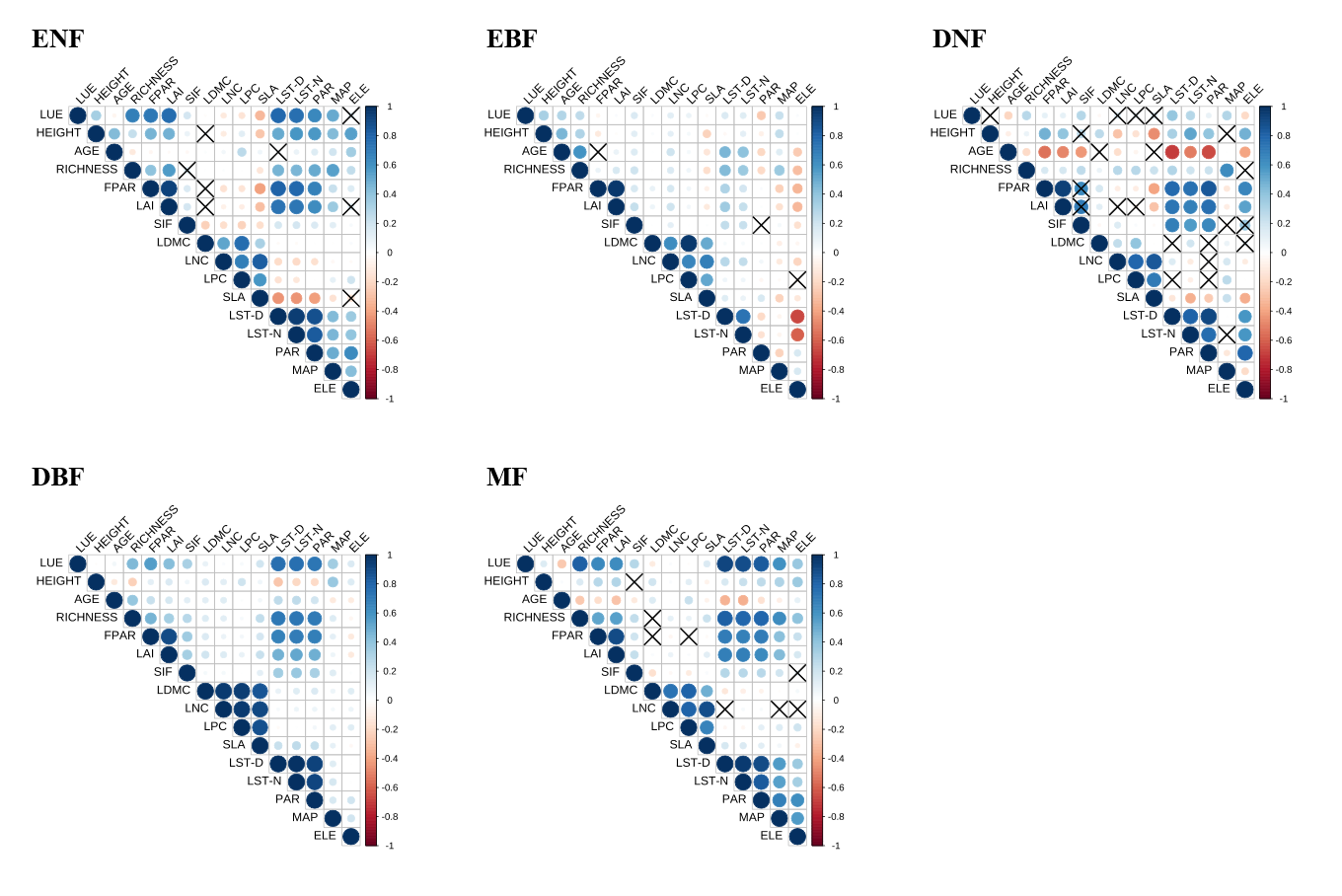

Figure S2 Correlation matrix plot of LUE and multiple plant traits and climatic conditions for 5 forest types. × means not significant.

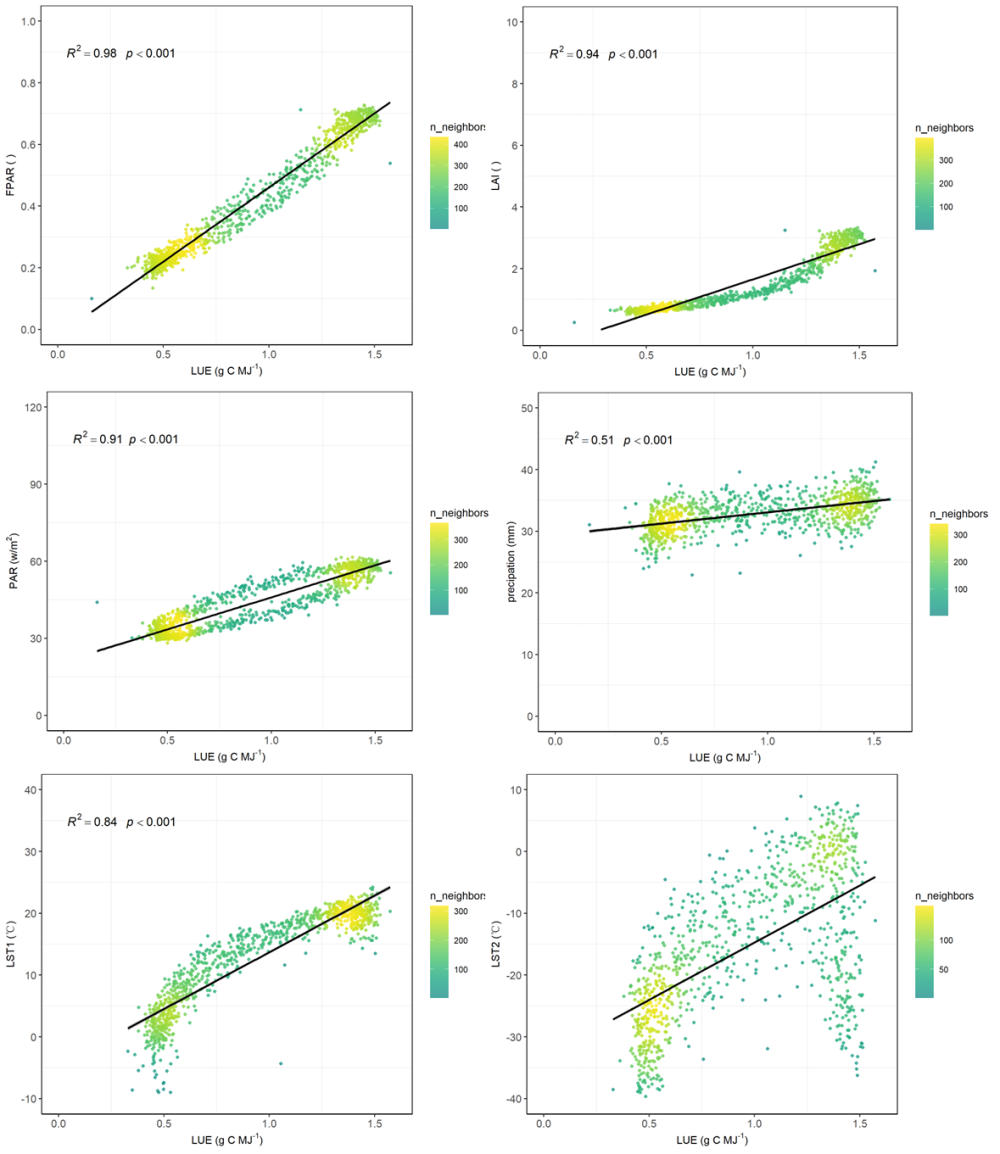

Figure S3 Scatter plot of LUE and multiple plant functional traits and environmental conditions in temporal analysis.

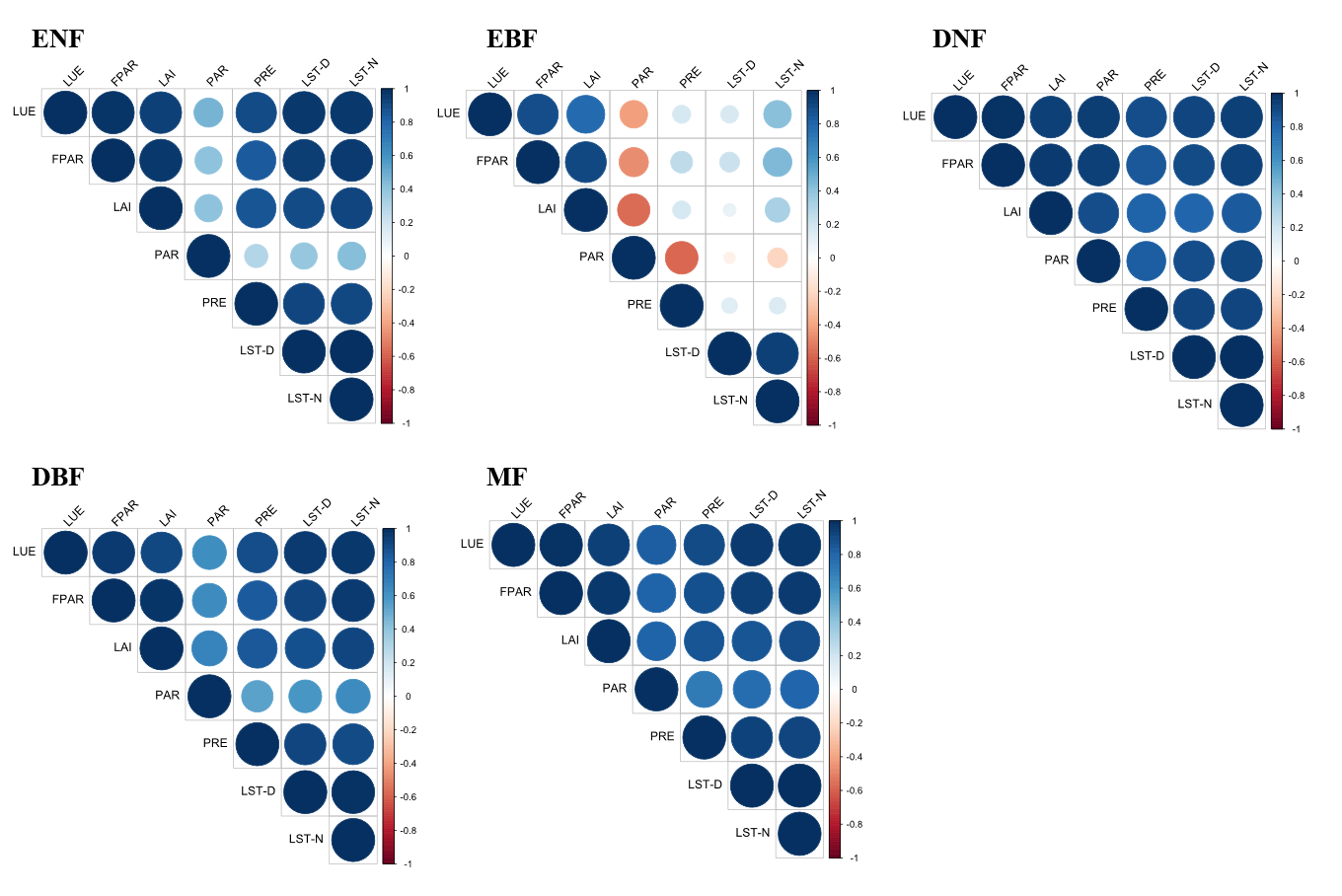

Figure S4 Correlation matrix plot of LUE and multiple plant traits and climatic conditions for 5 forest types. × means not significant.
